# Elimination of GPI2 suppresses glycosylphosphatidylinositol GlcNAc transferase activity and alters GPI glycan modification in *Trypanosoma brucei*

**DOI:** 10.1101/2021.03.18.436003

**Authors:** Aurelio Jenni, Sebastian Knüsel, Rupa Nagar, Mattias Benninger, Robert Häner, Michael A. J. Ferguson, Isabel Roditi, Anant K. Menon, Peter Bütikofer

## Abstract

The biosynthesis of glycosylphosphatidylinositol (GPI) membrane protein anchors is initiated in the endoplasmic reticulum by transfer of GlcNAc from the sugar nucleotide UDP-GlcNAc to phosphatidylinositol. The reaction is catalyzed by GPI GlcNAc transferase, a multi-subunit complex comprising the catalytic subunit Gpi3/PIG-A, as well as at least five other subunits including the hydrophobic protein Gpi2 which is essential for activity in yeast and mammals, but whose function is not known. Here we exploited *Trypanosoma brucei* (Tb), an early diverging eukaryote and important model organism, to investigate the function of Gpi2. We generated trypanosomes that lack TbGPI2 and found that in TbGPI2-null parasites (i) GPI GlcNAc transferase activity is reduced but not lost, in contrast with the situation in yeast and human cells, (ii) the GPI GlcNAc transferase complex persists, but its architecture is affected, with loss of at least the TbGPI1 subunit, and (iii) the GPI anchors of the major surface proteins are underglycosylated when compared with their wild-type counterparts, indicating the importance of TbGPI2 for reactions that are expected to occur in the Golgi apparatus. Additionally, TbGPI2-null parasites were unable to perform social motility, a form of collective migration on agarose plates. Immunofluorescence microscopy localized TbGPI2 to the endoplasmic reticulum as expected, but also to the Golgi apparatus, suggesting that in addition to its expected function as a subunit of the GPI GlcNAc transferase complex, TbGPI2 may have an enigmatic non-canonical role in Golgi-localized GPI anchor modification in trypanosomes.

## Introduction

Roughly 1% of all proteins encoded by eukaryotic genomes are post-translationally modified at their C-terminus by glycosylphosphatidylinositol (GPI), a complex glycoglycerophospholipid that anchors the protein to the cell surface. Its core structure consists of ethanolamine-PO_4_-6Manα1-2Manα1-6Manα1-4GlcNα1-6*myo*-inositol-1-PO_4_-lipid, with the ethanolamine residue being linked to the C-terminus of the protein via an amide bond (1). The glycan core can be extensively modified with phosphoethanolamine residues, mono- and/or oligosaccharides depending on the protein and cell-type in question (2).

GPI anchoring occurs in the lumen of the endoplasmic reticulum (ER), but the biosynthesis of the glycolipid itself is initiated on the cytoplasmic side (3) by the addition of GlcNAc from UDP-GlcNAc to a *myo*-inositol-containing phospholipid, most commonly phosphatidylinositol (PI). In subsequent reactions, GlcNAc-PI is de-*N*-acetylated (4) to glucosamine (GlcN)-PI, which is then translocated across the ER membrane (5) by an unknown scramblase and modified by addition of mannosyl and phosphoethanolamine residues to generate a GPI anchor precursor appropriately situated for transfer to newly translocated proteins (1, 2, 6). Species- and cell type-specific modifications of the GPI core structure then occur in the ER, Golgi and during transport of GPIs and GPI-anchored proteins to the cell surface (7–10).

The synthesis of GlcNAc-PI is catalyzed by UDP-GlcNAc : PI α1-6 GlcNAc-transferase (henceforth GPI GlcNAc transferase), a multi-subunit, membrane-bound complex consisting of Gpi1/PIG-Q, Gpi2/PIG-C, Gpi3/PIG-A, Gpi15/PIG-H, Gpi19/PIG-P, Eri1/PIG-Y (nomenclature corresponding to yeast/mammals) (11–20); in mammalian cells, a seventh subunit - Dpm2 - has been reported (12). The multi-subunit nature of this enzyme is unexpected and enigmatic. While it is clear that Gpi3/PIG-A is the catalytic subunit (21, 22), the functions of the other subunits are not evident. We were intrigued by the Gpi2/PIG-C subunit, a highly hydrophobic membrane protein that is essential for GPI GlcNAc transferase activity in yeast and humans (11, 18, 23). It has been speculated that Gpi2/PIG-C might play a role in recruiting the hydrophobic lipid substrate, PI, to the GPI GlcNAc transferase complex and/or maintaining the architecture of the transferase complex. To explore these possibilities we turned to the parasite causing human sleeping sickness, *Trypanosoma brucei*, which offers a number of genetic and biochemical advantages to study GPI anchoring, notably that GPI biosynthesis is not essential for the survival of *T. brucei* procyclic forms in culture (24, 25), conveniently allowing manipulation of the GPI pathway without compromising cell viability. Historically, the high abundance of GPIs and GPI-anchored proteins in trypanosomes made it possible to delineate the first complete structure of a GPI anchor in *T. brucei* bloodstream forms (26) and the corresponding anchors in insect stage (procyclic) forms (27–29), and to elucidate the reaction sequences leading to their synthesis (30–34). Notably, the GPI anchors in *T. brucei* procyclic forms are among the most complex GPI structures identified to date, with unusually large side chains consisting of characteristic poly-disperse-branched *N*-acetyllactosamine (Galβ1-4GlcNAc) and lacto-*N*-biose (Galβ1-3GlcNAc) units capped with sialic acid residues (27, 28). Several enzymes involved in GPI side chain modification in *T. brucei* have been identified and characterized (7, 35–37).

The core subunits of the GPI GlcNAc transferase complex have been identified in *T. brucei* by bioinformatics (38) and quantitative proteomics (39): TbGPI1 (Tb927.3.4570), TbGPI2 (Tb927.10.6140), TbGPI3 (Tb927.2.1780), TbGPI15 (Tb927.5.3680), TbGPI19 (Tb927.10.10110), and TbERI1 (Tb927.4.780) (TbDPM2 (Tb927.9.6440) is also listed in the *T. brucei* genome but this may be a mis-annotation as the trypanosome dolichol phosphate mannose synthase, like its yeast counterpart, comprises a single protein, TbDPM1 (40)). To explore the role of TbGPI2, we deleted the gene in *T. brucei* procyclic forms and characterized the knockout cells (TbGPI2-KO) using a variety of biochemical readouts. The results of our analyses were unexpected at multiple levels and showed that GPI GlcNAc transferase activity is reduced but not lost in TbGPI2-KO parasites, and that whereas the GPI GlcNAc transferase complex persists, its architecture is affected, with loss of at least the TbGPI1 subunit. Unexpectedly, we found that GPI anchors of the major surface glycoproteins are underglycosylated in the absence of TbGPI2, indicating the importance of this protein for reactions that are expected to occur in the Golgi apparatus and suggesting that TbGPI2 may possess a hitherto unknown non-canonical function in regulating GPI side chain modification in the Golgi apparatus.

## Results and Discussion

### TbGPI2 is not required for growth of *T. brucei* procyclic forms

To investigate the role of TbGPI2 in GPI biosynthesis in *T. brucei*, we used CRISPR/Cas9 to replace both alleles of TbGPI2 with antibiotic resistance cassettes in procyclic form parasites. One viable clone was obtained and replacement of both TbGPI2 alleles with drug resistance cassettes was verified by PCR (Fig. S1A) and Southern blotting (Fig. S2A). Loss of TbGPI2 mRNA was verified by Northern blotting (Fig. S2B). The TbGPI2 knock-out (TbGPI2-KO) parasites grew more slowly than the isogenic parental strain, with a doubling time of ∼11 h compared to ∼9.4 h for parental cells (Fig. 1A). Slower growth of GPI-deficient procyclic cells has been reported previously in some instances, for example after knocking out TbGPI13 or TbGPI10 (24, 41), but not TbGPI12 (25). Growth was restored by expressing an ectopic copy of TbGPI2 (TbGPI2-HA, bearing a C-terminal 3x HA tag) in the TbGPI2-KO parasites (Fig. 1A; integration of the ectopic copy was verified by PCR (Fig. S1B) and expression of HA-tagged TbGPI2 by SDS-PAGE/immunoblotting (Fig. S1C)).

**FIGURE 1:**
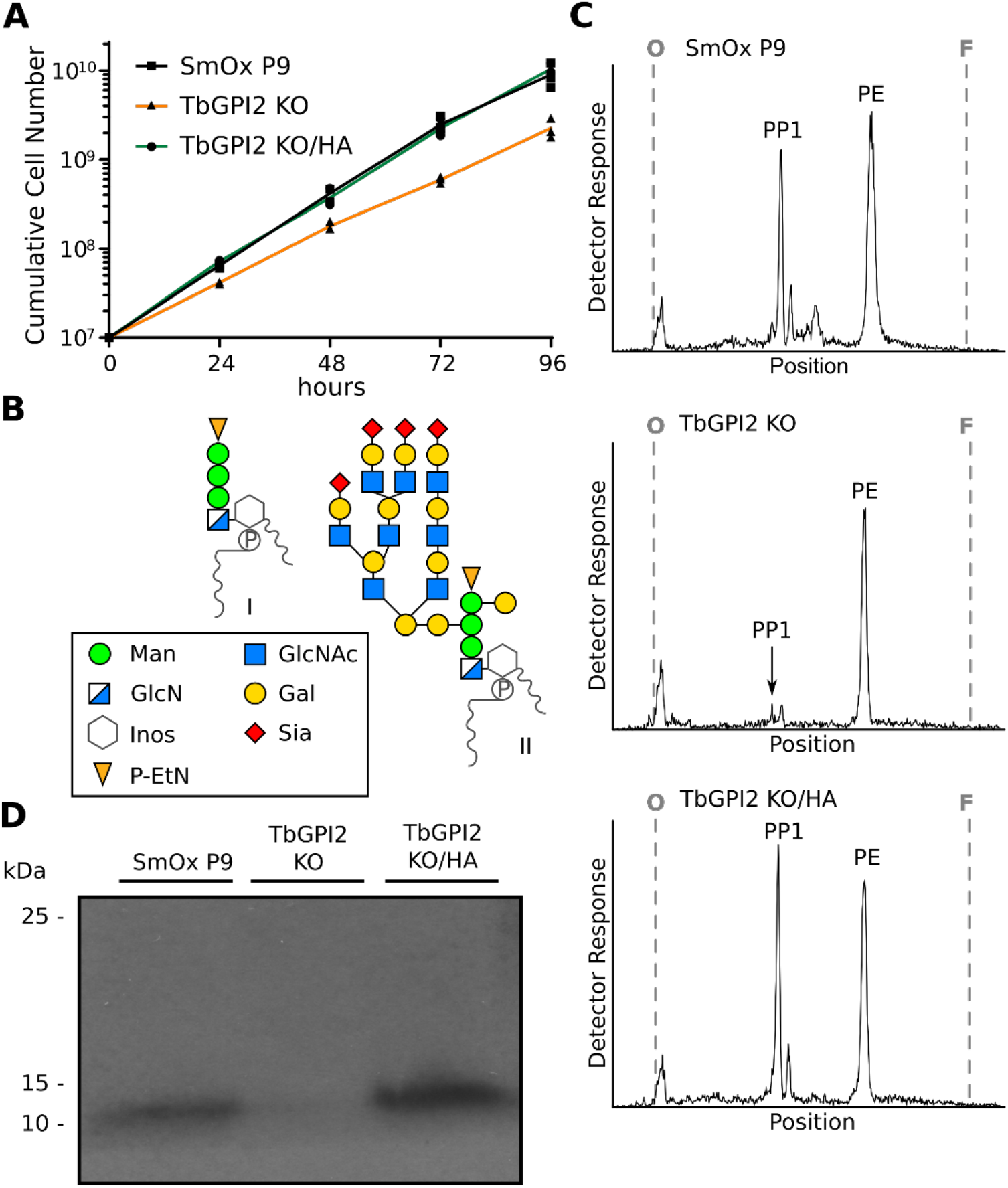
Characterization of TbGPI2-KO parasites. A. Growth of *T. brucei* SmOx P9 (black line), TbGPI2-KO (red line) and TbGPI2-KO/HA (green line) procyclic forms cultured under identical conditions. Data are from three independent experiments. B. Schematic structures of GPI precursor PP1 (I) and free GPIs (II, taken from the procyclin GPI anchor). C, D. *T. brucei* SmOx P9, TbGPI2-KO and TbGPI2-KO/HA were cultured for 16 h in the presence of [^3^H]-ethanolamine and subjected to a sequential extraction protocol. GPI precursors and free GPIs were analyzed by TLC and radioisotope scanning (C; O, site of sample application; F, solvent front; the extracts contain residual amounts of [^3^H]-ethanolamine-labeled phosphatidylethanolamine (PE)), and SDS-PAGE followed by fluorography (D; molecular mass markers are indicated), respectively.

We next tested the ability of TbGPI2-KO parasites to synthesize GPIs. Trypanosomes were metabolically labeled with [^3^H]-ethanolamine, and GPI precursors and free GPIs (Fig. 1B) were sequentially extracted and analyzed by TLC. Unexpectedly, extracts containing GPI precursors revealed the presence of small amounts of PP1 (<20% of that in parental cells) (Fig. 1C), suggesting residual GPI GlcNAc transferase activity in TbGPI2-KO parasites. This result contrasts with findings from yeast (11) and human (18, 42) cells, where disruption of ScGPI2 and PIG-C, respectively, results in total loss of GPI GlcNAc transferase activity. In addition, we found that the levels of free GPIs were decreased in TbGPI2-KO parasites compared to parental cells (<30% of control values; Fig. 1D). Expression of TbGPI2-HA in the TbGPI2-KO background completely restored both PP1 precursor and free GPI levels (Fig. 1C, D), indicating that TbGPI2-HA is functional and that the residual activity in TbGPI2-KO cells is not due to some form of adaptation in culture. We conclude that TbGPI2 has an important yet non-essential contribution to the activity of GPI GlcNAc transferase in *T. brucei*, such that a low level of GPI biosynthesis persists even in the absence of TbGPI2.

### TbGPI2-KO-derived membranes are able to synthesize GlcNAc-PI

To quantify GPI GlcNAc transferase activity in TbGPI2-KO cells, we used a cell-free assay in which crude membranes are tested for their ability to generate [^3^H]GlcNAc-PI from UDP-[^3^H]GlcNAc and endogenous PI (30, 31, 33). TLC analyses of lipid extracts from such assays showed that whereas both parental (Fig. 2A) and TbGPI2-KO (Fig. 2B)-derived membranes generated [^3^H]GlcNAc-PI and the product of the subsequent reaction, [^3^H]GlcN-PI, the reaction proceeded more slowly in TbGPI2-KO-derived membranes compared to membranes from parental cells (Fig. 2C, D).

**FIGURE 2:**
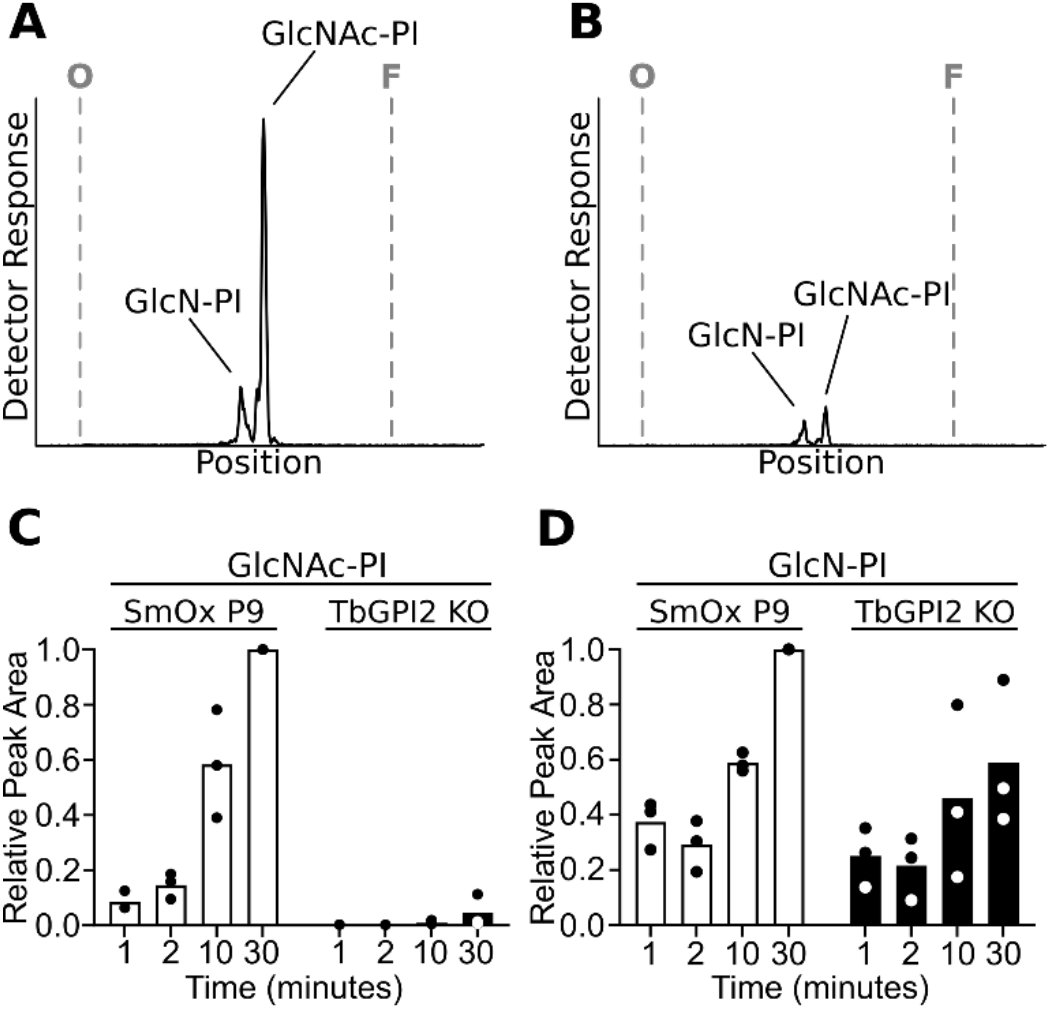
GPI GlcNAc transferase activity assayed in TbGPI2-KO membranes. A, B. Membranes from *T. brucei* SmOx P9 (A) and TbGPI2-KO (B) procyclic forms were incubated with UDP-[^3^H]GlcNAc for 30 min and lipids were extracted and analyzed by TLC and radioisotope scanning. O, site of sample application; F, solvent front. C, D. Quantification of time-dependent formation of [^3^H]GlcNAc-PI (C) and [^3^H]GlcN-PI (D) using peak area from the chromatograms as read-out. Data from three independent experiments are shown, normalized to parental SmOx P9 cells, 30 min incubation time.

### TbGPI2-KO cells synthesize GPI-anchored procyclins with reduced apparent mass

Since TbGPI2-KO parasites have a low level of GPI GlcNAc transferase activity and synthesize PP1, albeit in low amounts, we next investigated whether they also synthesize the stage-specific GPI-anchored procyclins EP (43) and GPEET (44). Procyclins can be extracted from cells with 9% (v/v) n-butanol in water (27, 44) and they migrate on SDS-PAGE as a broad band at 22-32 kDa for GPEET, and at around 42 kD for EP, which is a relatively minor GPI-anchored protein in early procyclic forms (44–46). TbGPI2-KO and isogenic parental cells were metabolically labeled with [^3^H]-ethanolamine, and probed for radiolabeled procyclins by SDS-PAGE and fluorography. Parental cells (SmOx P9) showed a strong radiolabeled GPEET band, as well as a weak band corresponding to EP procyclin (Fig. 3A), and as expected (41), no labeled procyclins were detected in a control sample of *T. brucei* procyclic forms that lack TbGPI13, the enzyme that adds phosphoethanolamine to the third mannose of the GPI anchor. Interestingly, TbGPI2-KO cells showed a radiolabeled band with a lower apparent mass than GPEET procyclin in parental cells. This result suggests that GPI anchoring of GPEET occurs in TbGPI2-KO cells, but that some aspect of GPEET maturation is disrupted resulting in a lower molecular weight form. Of note, we detected a radiolabeled 55 kDa protein in all three cell lines corresponding to ethanolamine phosphoglycerol-modified eukaryotic elongation factor 1a (eEF1A) (47) - the appearance of this band serves as a control for [^3^H]-ethanolamine labeling in our experiments.

**FIGURE 3:**
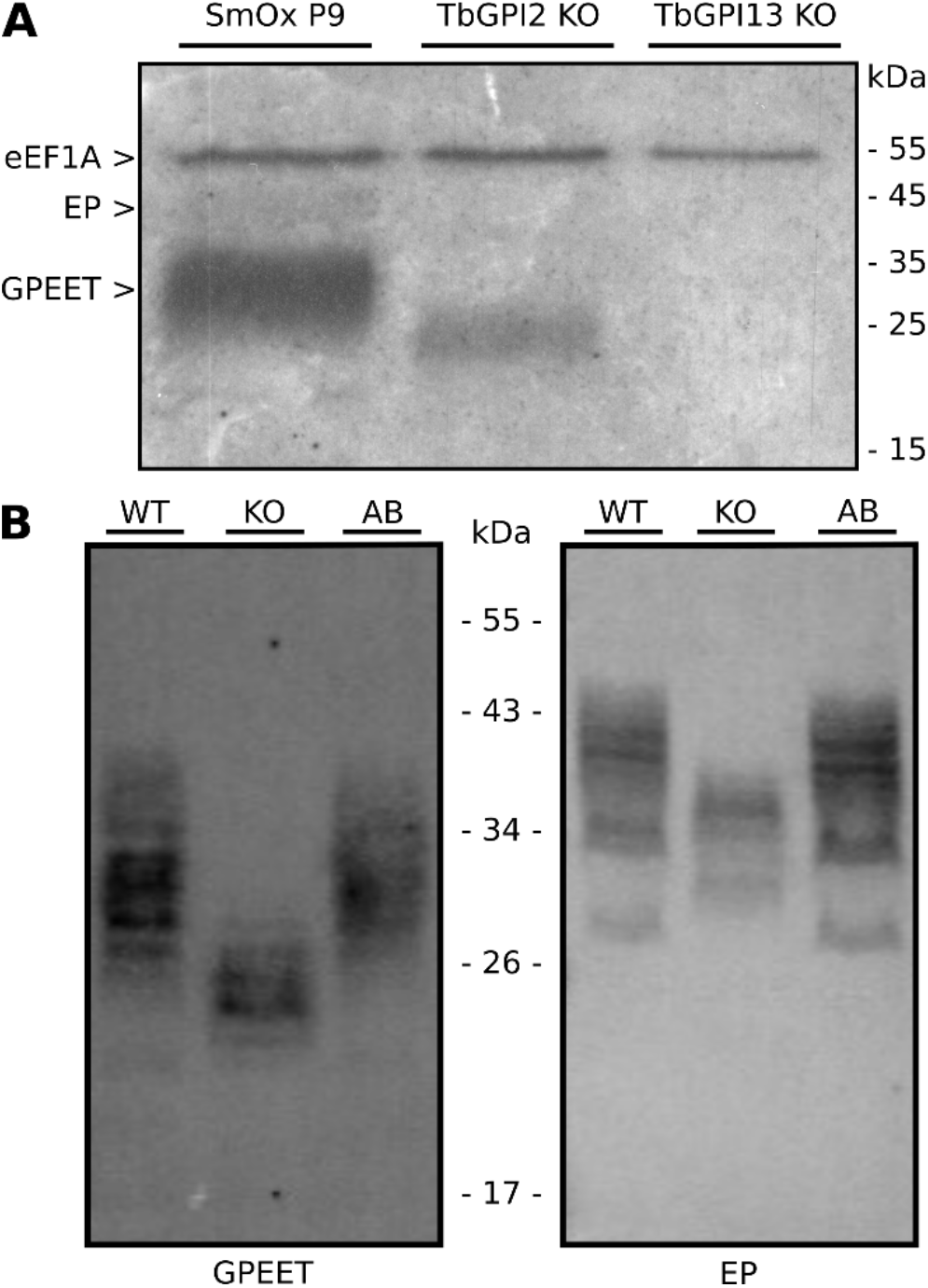
Analyses of GPEET and EP procyclin in TbGPI2-KO cells. A. *T. brucei* SmOx P9, TbGPI2-KO and TbGPI13-KO parasites were grown for 16 h in the presence of [^3^H]-ethanolamine. Proteins were analyzed by SDS-PAGE followed by fluorography. B. *T. brucei* SmOx P9 (WT), TbGPI2-KO (KO) and TbGPI2-KO/HA (AB) parasites were cultured under standard conditions and proteins were analyzed by SDS-PAGE followed by immunoblotting using α-GPEET 5H3 (left panel) or α-EP247 (right panel) antibodies. Molecular mass markers are indicated in the margins.

To extend the results of our radiolabeling experiments we probed for GPEET and EP in TbGPI2-KO parasites by immunoblotting using specific antibodies. The results show that TbGPI2-KO cells express both GPEET and EP (Fig. 3B) and that both procyclins had distinctly lower apparent masses in TbGPI2-KO compared to parental parasites. Again, expression of HA-TbGPI2 in the TbGPI2-KO background completely restored the parental phenotypes (Fig. 3B).

Previous studies showed that disruption of the GPI biosynthesis pathway may lead to retention and accumulation of normally GPI-anchored procyclins inside the cell (48). Although TbGPI2-KO cells retain the ability to synthesize GPI-anchored procyclins as shown above, we considered whether these proteins are indeed trafficked to the cell surface. Using immunofluorescence microscopy we found a typical surface staining pattern for both EP and GPEET procyclin in TbGPI2-KO parasites that was indistinguishable from that of parental cells (Fig. 4A). Quantitative analysis by flow cytometry revealed a slight decrease in surface-localized EP procyclin while the levels of surface-localized GPEET were unchanged (Fig. 4B). The parental phenotype was completely restored on expressing HA-TbGPI2 in TbGPI2-KO parasites (Fig. 4B). Thus, TbGPI2-KO cells synthesize non-native, GPI-anchored procyclins of a lower molecular weight that nonetheless are trafficked to the cell surface.

**FIGURE 4:**
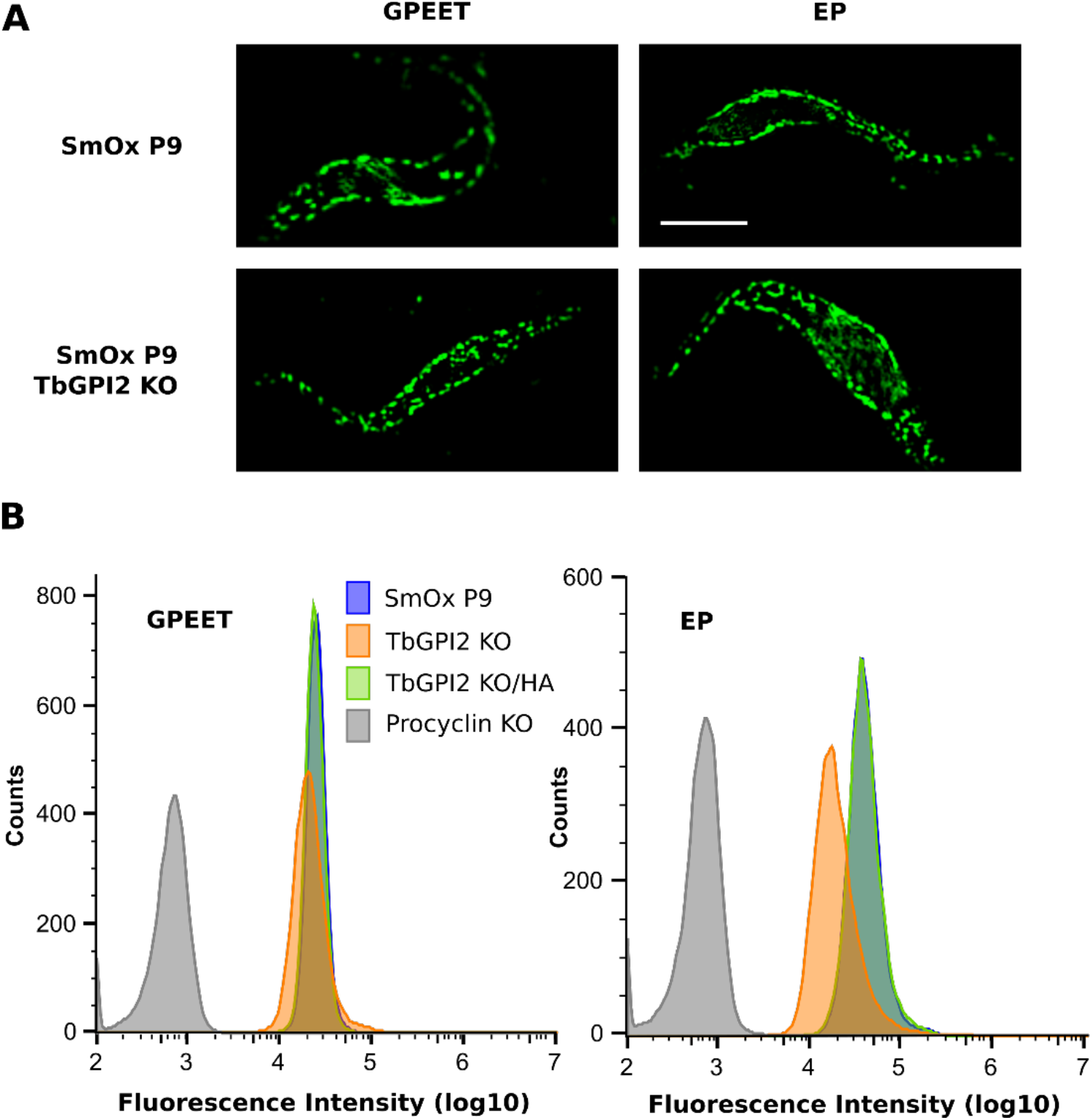
Surface localization of EP and GPEET procyclins. A. *T. brucei* SmOx P9 (top panels) and TbGPI2-KO (bottom panels) parasites were fixed with paraformaldehyde, and procyclins were visualized by fluorescence microscopy using α-GPEET 5H3 (left panels) or α-EP 247 (right panels) antibodies in combination with the corresponding fluorescent secondary antibodies. Scale bar, 5 µm. B. *T. brucei* SmOx P9 (blue), TbGPI2-KO (orange) and TbGPI2-KO/HA (green) parasites were labeled as in A, and surface labeling of GPEET and EP was quantified by flow cytometry. Note that the traces for SmOx P9 and TbGPI2-KO cells almost completely overlap. *T. brucei* procyclin null parasites (Procyclin KO) were used as a negative control.

### GPI anchors in TbGPI2-KO parasites are underglycosylated

We hypothesized that the non-native procyclin structures seen in TbGPI2-KO cells are due to under-glycosylation of the GPI anchor. To investigate this, we purified procyclins from parental, TbGPI2-KO mutant and TbGPI2-KO mutants expressing TbGPI2-HA (add-back cells), and subjected them to aqueous hydrofluoric acid dephosphorylation, a process that liberates GPI anchor glycans from the procyclin polypeptide and *lyso*-phosphatidic acid lipid components of the PI moiety. The released GPI glycans were subsequently permethylated, a procedure that methylates all free hydroxyl groups and converts the amine group of the glucosamine residue to a positively charged trimethyl quaternary ammonium ion, and which also removes the fatty acid from the inositol ring of GPI glycans. The permethylated GPI glycan fractions were analysed by positive ion ES-MS (7, 8). The TbGPI2-KO mutant sample showed the presence of a series of triply charged [M + 2Na]^3+^ precursor ions in MS^1^ (Fig. S3) corresponding to GPI glycans with Hex-HexNAc repeats with and without sialic acid (7, 8), similar to the samples from the parental and add-back cell lines. The triply charged [M + 2Na]^3+^ ions and their proposed molecular compositions for the TbGPI2-KO sample are shown in (Table S1). The smallest and largest GPI glycans observed in the TbGPI2-KO mutant sample were (Gal_5_GlcNAc_2_)Man_3_GlcN-Ino and (Gal_11_GlcNAc_8_SA_1_)Man_3_GlcN-Ino, respectively.

The identities of triply charged [M + 2Na]^3+^ GPI glycan ions were confirmed by MS^2^ and MS^3^ analyses. For example, the MS^2^ spectra of triply charged [M + 2Na]^3+^ ions at *m/z* 888.45 (consistent with compositions of permethylated (Gal_5_GlcNAc_2_)Man_3_GlcN(Me_3_)^+^Ino in samples from parental, add-back and TbGPI2-KO cells all produced intense doubly-charged [M + 2Na]^2+^ fragment ions at *m/z* 1186.56. The latter arises from the characteristic facile elimination of the inositol residue and quaternary amine group (Fig. S4) (7, 8). On further fragmentation (MS^3^) these *m/z* 1186.56 ions all produced similar product ion spectra consistent with the structure proposed (Fig. 5). These spectra strongly support the presence of GPI anchors on procyclins produced by TbGPI2-KO parasites, as well as by the parental and TbGPI2 add back cells as expected.

**FIGURE 5:**
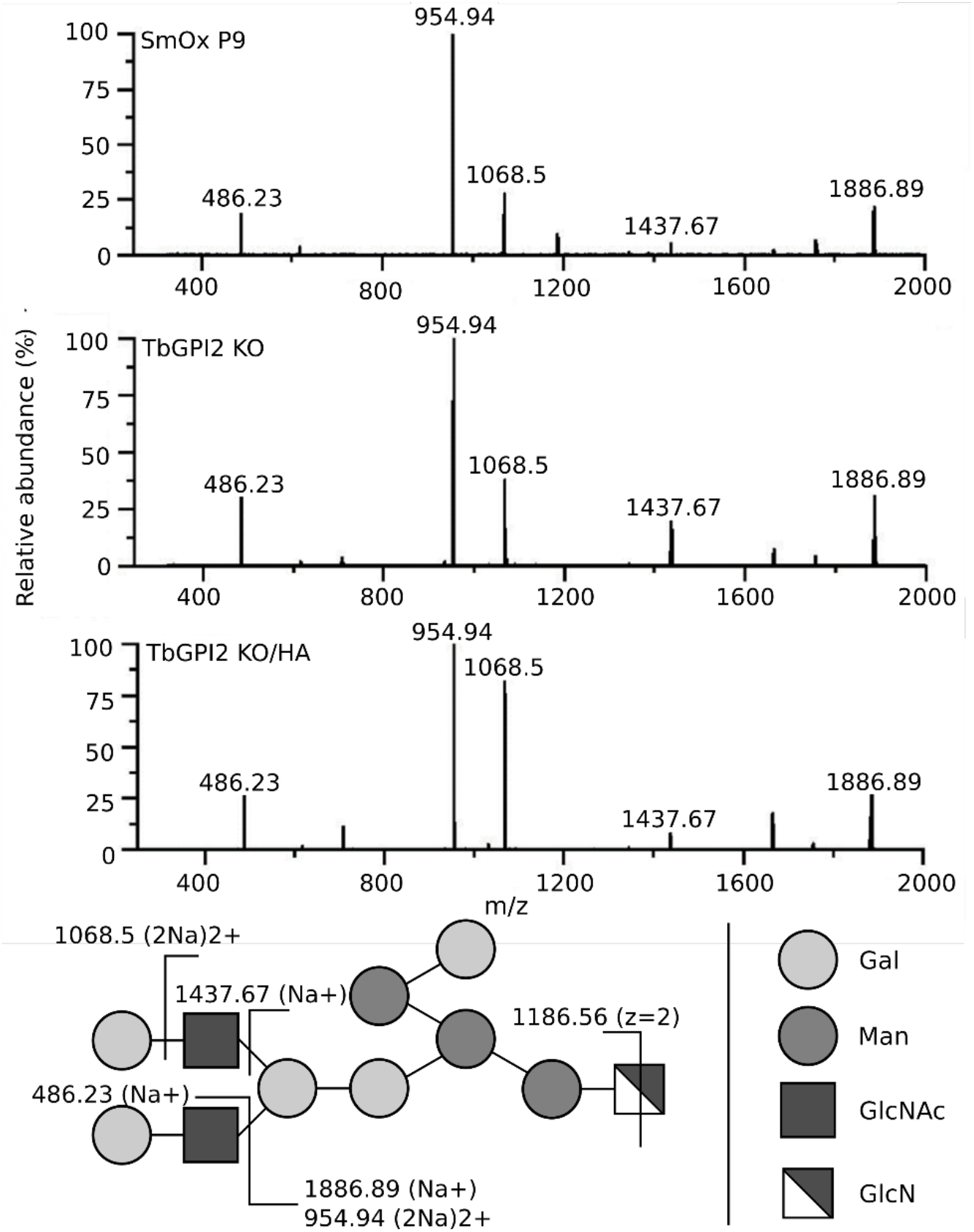
MS^3^ product ion spectra of permethylated GPI-glycans in TbGPI2-KO and control parasites. Permethylated GPI glycans from parental (top), TbGPI2-KO (middle) and TbGPI2-KO/HA (bottom) mutant parasites were analyzed by positive ion ES-MS. Triply charged [M + 2Na]^3+^ ions observed at *m/z* 888.45 for each sample were fragmented (MS^2^) generating a doubly charged product ion at *m/z* 1186.56 (see Fig. S4). This ion was further fragmented (MS^3^) to generate the product ion spectra shown. Assignments of the major product ions are indicated on the inset diagram.

Interestingly, the triply charged GPI glycan ions were more intense in the MS^1^ spectrum of the TbGPI2-KO mutant sample than in the parental and TbGPI2-KO/HA add back samples. We interpret this as being consistent with the TbGPI2-KO mutant GPI glycans being smaller than those of the wild type and TbGPI2-KO/HA add back, since the larger the molecular species the harder they are to ionise and observe in MS^1^. Our results suggest, therefore, that the ∼10 kDa lower molecular weight of GPEET seen in SDS-PAGE fluorograms of [^3^H]-ethanolamine labeled cells is due to a reduction in the number of LacNAc and/or lacto-N-biose repeats (Fig. 3A).

### TbGPI2 depletion leads to defects in social motility and growth on semisolid surfaces

We previously showed that a *T. brucei* mutant with a perturbation in *N*-linked glycosylation and GPI glycosylation (49) was impaired in its ability to perform social motility (SoMo), a form of collective migration on agarose plates, as well as to colonise the tsetse fly vector (50). We therefore examined the effect of the TbGPI2 deletion on SoMo. Our results reveal that TbGPI2-KO cells showed essentially no SoMo (Fig. 6A), and that the parental phenotype could be restored by expressing TbGPI2-HA (Fig. 6A). In addition, we noticed morphological abnormalities (Fig. 6B, upper panels) and decreased viability (Fig. 6C) of TbGPI2-KO parasites cultivated on agar plates compared to parental cells (Fig. 6B, upper panels). In liquid culture morphological abnormalities were only observed in a small fraction of cells (<5%).

**FIGURE 6:**
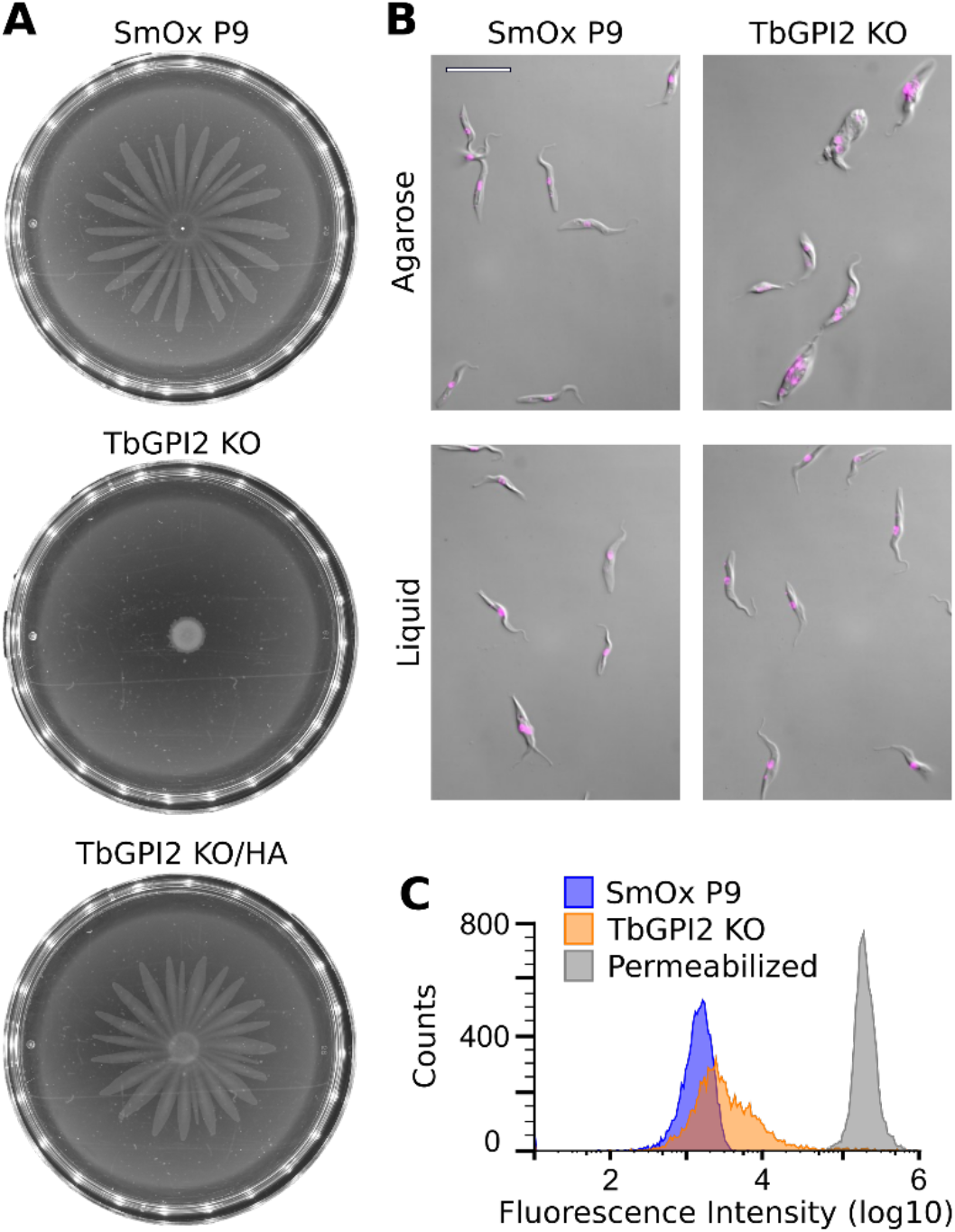
Behaviour of TbGPI2-KO trypanosomes grown on agarose plates. A. *T. brucei* SmOx P9 (top), TbGPI2-KO (middle) and TbGPI2-KO/HA (bottom) parasites were inoculated on agarose plates. The plates were photographed after 4 days of incubation at 27 °C. B. Parasites grown on agarose plates for 2 days and subsequently washed off (top) or in liquid culture for 2 days (bottom) were stained with Hoechst dye and examined by light microscopy. Scale bar, 20 µm. C. Parasites grown as in panel A were stained with propidium iodide and subjected to flow cytometry analysis.

Previous work indicated that procyclin null mutants are capable of SoMo (51); as these mutants compensated for the lack of procyclins by expressing free GPIs on their surfaces (52, 53), it appeared likely that the GPI moieties themselves might be sufficient for SoMo. This observation is consistent with our findings that mutants lacking TbRft1 (49) and TbGPI2 (this study) produce truncated GPI anchors and have fewer free GPIs. Thus the GPIs in these mutants are inadequate in terms of supporting SoMo despite the cells having normal levels of surface procyclins.

There are several parallels between SoMo and the swarming motility of bacteria on semi-solid surfaces. Cells face a number of challenges: they need to extract water from the surface in order to remain hydrated, and they must overcome friction and surface tension in order to move. Bacteria accomplish this by producing (lipo)polysaccharides and surfactants such as glycolipids or lipidated peptides (54–57). It is conceivable that GPIs act as lubricants facilitating movement, and that the glycocalyx of GPI-anchored proteins or free GPIs protects cells against dehydration.

### The GPI GlcNAc transferase complex is affected by the absence of TbGPI2

To study the effects of the absence of TbGPI2 on the GPI GlcNAc transferase complex, we selected TbGPI1, a multi-spanning membrane protein subunit of the complex, as a reporter. We epitope-tagged TbGPI1 in *T. brucei* SmOx P9 and TbGPI2-KO parasites and compared its expression level and inclusion in the GPI GlcNAc transferase complex in the two cell lines. Analysis by SDS-PAGE and immunoblotting showed that expression of cMyc-tagged TbGPI1 is unaffected by the absence of TbGPI2 (Fig. 7A). Native PAGE revealed that TbGPI1 from SmOx P9 parasites migrates as a broad band between the 242 kDa and 720 kDa molecular mass markers (Fig. 7B), reflecting its association with the GPI GlcNAc transferase complex. In a previous report (39), a GPI GlcNAc transferase complex isolated from *T. brucei* bloodstream by pull-down of cMyc-tagged TbGPI3 was seen to run closer to the 242 kDa marker. The reason for the difference on native PAGE between our data and the published report is not clear, but could be due to heterogeneity of the complexes revealed by using different baits for pull-down (TbGPI3 versus TbGPI1), or differences in the architecture of the complexes in bloodstream versus procyclic trypanosomes. Despite its comparable expression level in parental SmOx P9 and TbGPI2-KO cells, TbGPI1 was detected at much lower levels in native PAGE analysis of extracts from the TbGPI2-KO parasites (Fig. 7B), and appeared at the higher end of the molecular mass spectrum (480-720 kDa) seen for native GPI GlcNAc transferase. This result suggests that in the absence of TbGPI2, TbGPI1 is poorly recruited into the GPI GlcNAc transferase complex, and that complexes that do retain TbGPI1 run at a higher apparent molecular mass.

**FIGURE 7:**
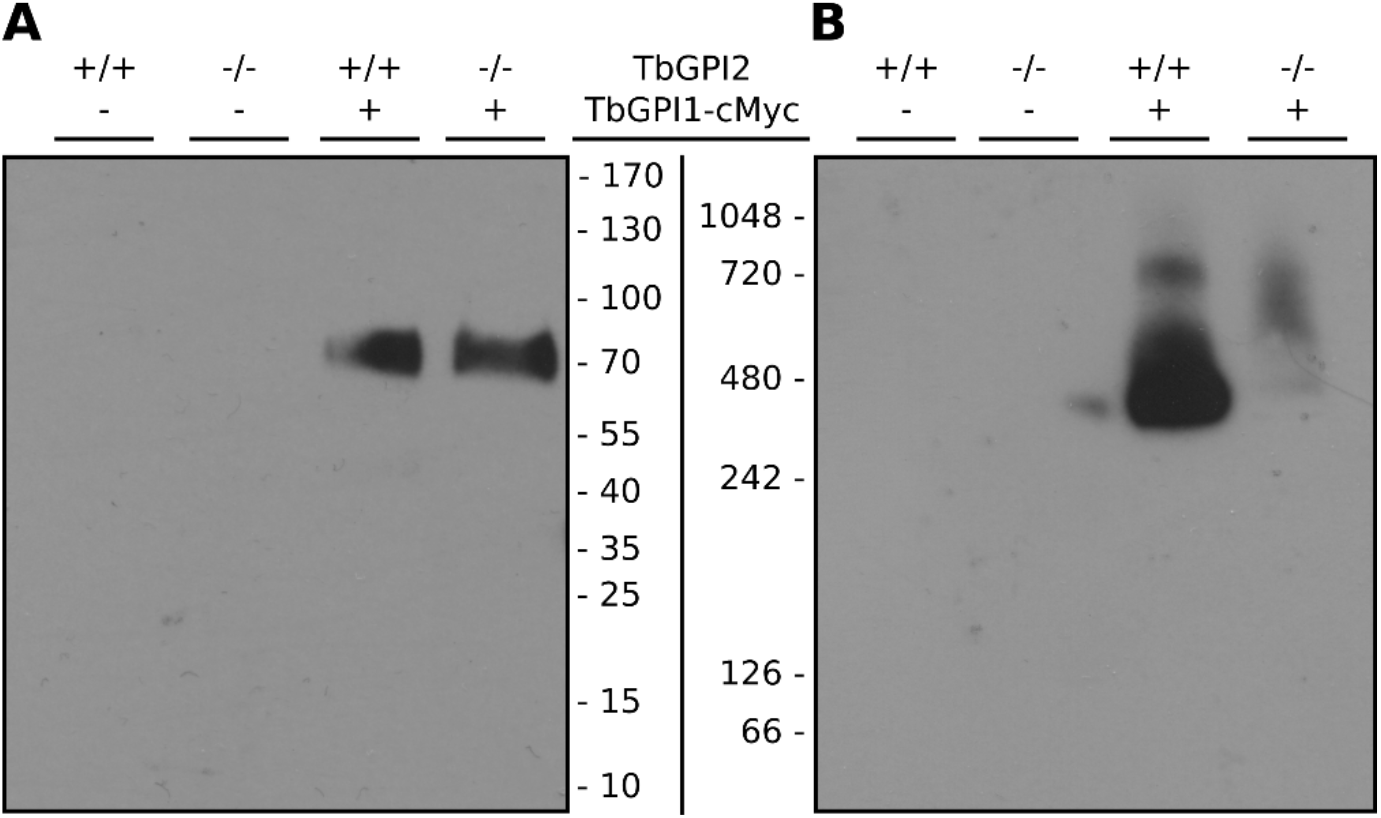
GlcNAc transferase complex in TbGPI2-KO trypanosomes. *T. brucei* SmOx P9 (+/+) and TbGPI2-KO (−/−) parental parasites (−), and SmOx P9 (+/+) and TbGPI2-KO (−/−) parasites expressing cMyc-tagged TbGPI1 (+), were immunoprecipitated from cell extracts prepared under non-denaturing conditions and analyzed by SDS-PAGE (panel A) and native PAGE (panel B). TbGPI1 was detected by immunoblotting using anti-cMyc antibody. The lanes contain identical cell equivalents. Molecular mass markers are indicated in kDa in the margins.

### TbGPI2 partially localizes to the Golgi apparatus

The phenotypic profile of TbGPI2-KO cells is reminiscent of some features of TbRft1-KO trypanosomes, specifically GPI underglycosylation (49) and SoMo defects (50). Immunofluorescence microscopy revealed that TbRft1 was localized to both the ER and the Golgi apparatus, hinting at a possible explanation for its role in regulating Golgi-localized GPI glycosylation (49). Because of the link between TbGPI2 and GPI glycosylation, we considered that TbGPI2 may also be localized to both the ER and Golgi apparatus.

Immunofluorescence microscopy of TbGPI2-HA-expressing cells showed extensive co-staining with the ER resident lumenal chaperone TbBiP (Fig. 8), indicating the protein was localized to the ER as expected. However, in >67% of cells (n=122), TbGPI2-HA was found to also co-localize with the Golgi-resident protein TbGRASP (58). We previously showed that (i) this level of co-localization with a Golgi marker is highly significant and not a random occurrence as it is detected in fewer than 35% of cells expressing an epitope-tagged version of the ER resident protein TbEMC3 (Tb927.10.4760) (49), and (ii) it is unlikely to be a result of saturation of the retention system for ER resident proteins (49). We conclude that TbGPI2 is localized to both the ER and Golgi apparatus.

**FIGURE 8:**
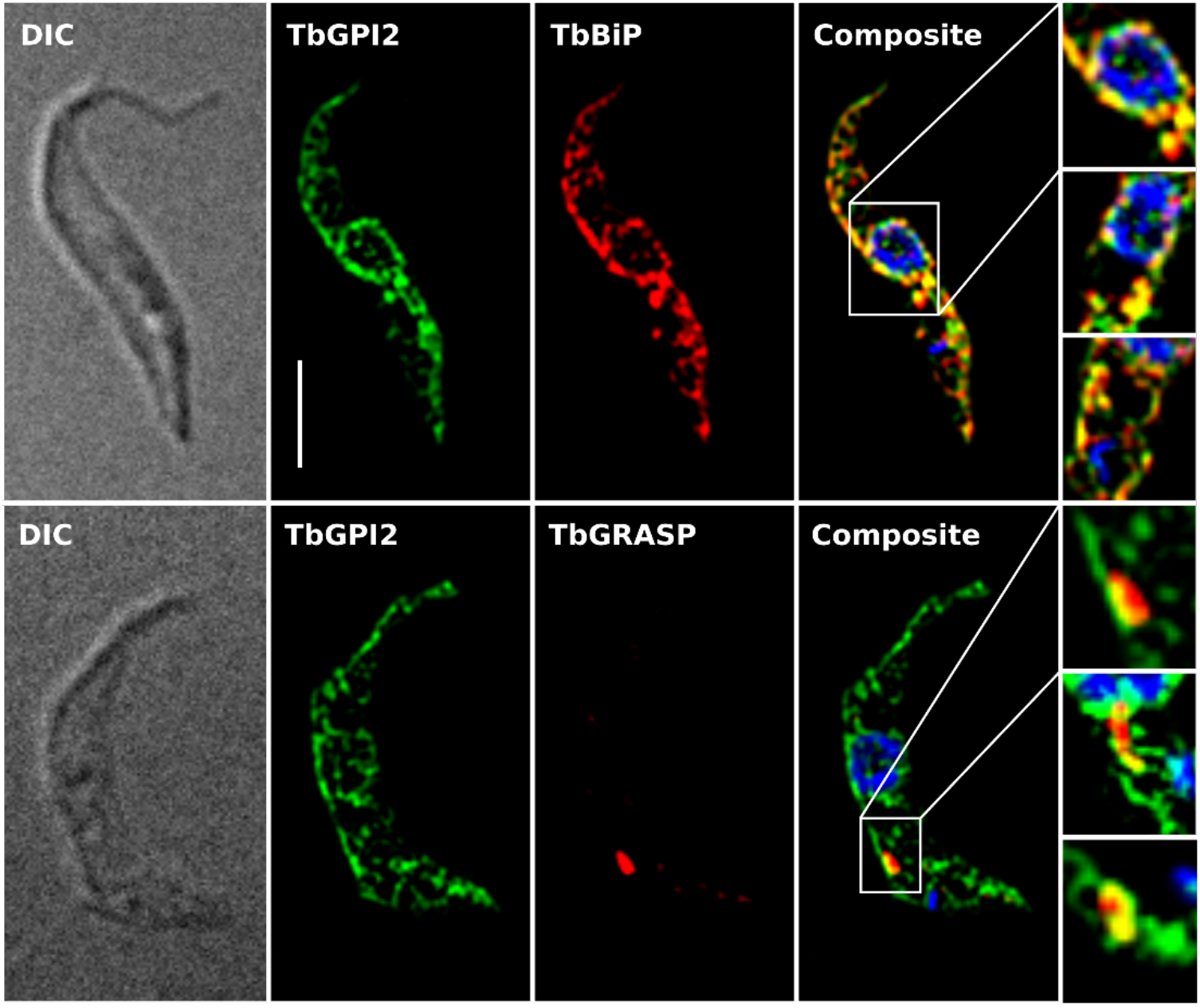
Localization of TbGPI2 in procyclic form parasites. Functional TbGPI2-HA (green) was visualized by indirect immunofluorescence along with the ER marker protein TbBiP or the Golgi-resident protein TbGRASP (red). DNA was stained with DAPI (blue). Co-localization of TbGPI2 and TbBiP/TbGRASP is shown in yellow (composite). Areas of interest from multiple cells are shown in the enlargements on the right. Scale bar, 10 µm.

## Concluding remarks

Gpi2/PIG-C is an essential component of the GPI GlcNAc transferase complex in yeast (11, 23) and human (18) cells, but its precise function is not known. We now report that TbGPI2 is important but not essential for *T. brucei* GPI GlcNAc transferase activity, which persists in TbGPI2-KO parasites at a low level as seen by *in vitro* assays and metabolic labeling experiments. In the absence of TbGPI2, the architecture of the GPI GlcNAc transferase complex is compromised, losing most of its content of TbGPI1. This disruption suggests a central scaffolding role for TbGPI2 in recruiting/organizing other subunits of the complex, different from the organization of the yeast and mammalian complex (59) where Gpi1 was proposed to link Gpi3 and other subunits to Gpi2. More work needs to be done to sort out the arrangement and stoichiometry of the various subunits in the complex in order to understand their individual roles.

TbGPI2-KO trypanosomes are able to synthesize GPI-anchored proteins, but their GPI anchors are underglycosylated. This may underly their inability to perform SoMo as discussed above. As GPI glycosylation largely occurs in the Golgi apparatus (7, 8), it is possible that the Golgi pool of TbGPI2 that we observe may directly or indirectly influence the extent to which GPI side chains are extended. Perhaps TbGPI2 interacts with the enzymes responsible for GPI glycosylation, thereby affecting their function. This possibility could be tested in future studies aimed at detecting physical or genetic interaction partners of TbGPI2. It is also possible to rationalize GPI underglycosylation through a kinetics-based argument. Thus, loss of TbGPI2 decreases the rate of synthesis of GlcNAc-PI, which may lead to a reduction of the available amount of GPI anchors that can be exported to the cell surface. While procyclic form trypanosomes do not depend on a functioning GPI anchor pathway when cultured *in vitro* (24, 25), a growth defect is often observed when GPI anchor biosynthesis is disrupted (24, 41). Therefore, any increase of GPI-anchored proteins present on the cell surface is expected to reduce the growth defect and therefore increase the relative fitness. TbGPI2-KO trypanosomes which forgo certain GPI glycosylation steps in the Golgi are able to decrease time between GlcNAc-PI synthesis and export of the GPI anchor to the cell surface, thus increasing the amount of GPI-anchored proteins present on the plasma membrane at the cost of lowering the concentration of intermediate products of the pathway. This hypothesis is consistent with the decreased levels of PP1 and free GPIs in TbGPI2-KO parasites.

## Data availability

All data are contained within this manuscript.

## Supporting information

This article contains supporting information.

## Acknowledgements

We thank Monika Rauch and Jennifer Jelk for technical assistance during parts of the study. AJ thanks Sophie Stammherr for inspiring discussions. PB dedicates this paper to Annina Niederer (1953–2021) for her year-long support and encouragement.

## Materials and Methods

Unless otherwise stated, reagents were purchased from Merck KGaA (Darmstadt, Germany). Restriction enzymes were from ThermoFisher Scientific (Waltham, MA, USA). Tritium-labeled compounds were from American Radiolabeled Chemicals Inc. (St. Louis, MO, USA) and PCR reagents and restriction enzymes from Promega Corporation (Madison, WI, USA). Acrylamide mix was from National Diagnostics (Atlanta, GA, USA).

### Trypanosome Cultures

*T. brucei* SmOx P9 pTB011 procyclic forms (60) (henceforward termed SmOx P9) were maintained at 27 °C in SDM79 containing 10% (v/v) heat-inactivated fetal bovine serum, 160 μM hemin, 90 μM folic acid, 2 μg/ml puromycin and 5 μg/ml blasticidin. TbGPI2-KO, TbGPI2-KO/HA and *in situ*-tagged parasites were grown under the same conditions with additional selection antibiotics added (for TbGPI2-KO: 25 μg/ml hygromycin, 1 μg/ml G418; for TbGPI2-KO/HA: 25 μg/ml hygromycin, 1 μg/ml G418, 3.6 µg/ml phleomycin; for *in situ*: 3.6 µg/ml phleomycin).

### Generation of *T. brucei* TbGPI2-KO Parasites

TbGPI2-KO parasites were generated using CRISPR / CRISPR-associated protein 9 (Cas9) technique as described before (60). Briefly, two resistance gene cassettes were generated by PCR using primers 1 and 2 (Table S2) and template plasmid pPOTv6 (61) containing resistance genes for hygromycin and neomycin, respectively. The cassettes were flanked with homology sequences of 30 nt to replace both alleles of the target gene via homologous recombination. Two single-guide RNA templates containing a T7 polymerase promoter, a Cas9 binding site and a 20 nt targeting sequence were generated by PCR using primer pairs 3/5 and 4/5 (Table S2), respectively. All PCR were performed using the Expand High Fidelity PCR System (Roche Diagnostics GmbH, Mannheim, Germany) and primers were designed using the online tool at www.leishgedit.net. All PCR products were pooled and purified using the Wizard® SV Gel and PCR Clean-Up System (Promega). DNA (10 μg) was transfected into SmOx P9 cells using a 4D nucleofector system (Lonza Group AG, Basel, Switzerland) with program FI-115. After 24 h, selection antibiotics were added, the culture was diluted 1:25 and distributed into 24-well plates. TbGPI2 gene knock-out was verified by PCR using extracted gDNA from knock-out clones and primer pairs 6/7, 6/8, 6/9 and 6/10 (Table S2), as well as by Northern and Southern blotting (see below).

### Generation of *T. brucei* TbGPI2 Addback Parasites

TbGPI2 ORF was amplified from SmOx P9 gDNA using primer pairs 11/12 and 11/13 (Table S2), yielding untagged and HA-tagged constructs of TbGPI2 flanked by *Hind*III and *Xho*I restriction sites, which were then cloned into plasmid pMS1720RNAiBSF (62). Plasmids (10 μg) were then linearized with *Not*I and transfected into SmOx P9 TbGPI2-KO cells using a 4D nucleofector system (Lonza Group AG, Basel, Switzerland) with program FI-115. After 24 h, selection antibiotic was added, the culture was diluted 1:250 and distributed into 24-well plates. TbGPI2 gene addback was verified by PCR using extracted gDNA from addback clones and the following primer pairs: 6/7, 11/12 (Table S2). Protein expression was analyzed by SDS-PAGE followed by immunoblotting against the HA epitope.

### Northern and Southern blotting

TbGPI2 mRNA expression was assayed by Northern blotting as described before (43). Genomic DNA was isolated and digested with *Bgl*II or *Cla*I followed by Southern blotting as previously described (43). A radioactively labeled probe corresponding to the coding region of TbGPI2 was generated using a Megaprime DNA-labeling system (GE Healthcare) according to the manufacturer’s instructions. Signals were detected by exposure of the blots to a Phosphorimager screen (Amersham Biosciences, Amersham, UK) followed by scanning with a Typhoon FLA 7000 (GE Healthcare).

### In-Situ Tagging of TbGPI1

TbGPI1 was *in-situ* tagged in SmOx P9 and TbGPI2-KO cells using CRISPR/Cas9 technique as described above. Briefly, the resistance gene cassette was generated by PCR using primer pairs 14/15 as described above. The cassette consisted of a cMyc tag sequence and a phleomycin resistance gene and were flanked by homology sequences of 30 nt to insert between the last codon and the stop codon of the TbGPI1 by homologous recombination. A single-guide RNA template were generated by PCR using primer pairs 5/16 (TbGPI1) as described above. 10 μg of pooled and purified PCR were transfected into the cells as described above. 24 h after transfection, selection antibiotic was added, the cultures were diluted 1:100 (for SmOx P9) and 1:3 (for TbGPI2-KO) and distributed into 24-well plates. Protein expression was analyzed by SDS-PAGE followed by immunoblotting against the cMyc epitope.

### [^3^H]-Ethanolamine Labeling of GPI Precursors and GPI-Anchored Proteins

GPI precursors and GPI-anchored proteins were labeled and extracted as previously described (44). Briefly, 5 × 10^8^ trypanosomes were cultured for 16-18 h in the presence of 50 µCi of [^3^H]-ethanolamine. The cells were harvested by centrifugation and washed twice with Tris-buffered saline (TBS; 140 mM NaCl, 10 mM Tris, pH 7.4). Subsequently, phospholipids were extracted twice with chloroform/methanol 2:1 (v/v; CM fraction). GPI precursors and free GPIs (52, 63) were extracted three times with chloroform/methanol/water 10:10:3 (v/v/v; CMW fraction) and GPI-anchored proteins were extracted twice with 9 % butan-1-ol in water (v/v; BuOH fraction). The CMW fractions were further partitioned between butan-1-ol and water, yielding fractions CMW_but_ and CMW_aq_ containing GPI precursors and free GPIs, respectively. All fractions were pooled and dried and aliquots were used for liquid scintillation counting.

### Thin-Layer Chromatography

CMW_but_ or butanol extracts containing GPI precursors were resolved by TLC using Silica Gel 60 plates and chloroform/methanol/water (10:10:3; v/v/v) as the solvent system. The chromatograms were visualized using a Raytest Rita* radioactivity TLC analyser (Berthold Technologies, Regensburg, Switzerland).

### Protein Analysis

CMW_aq_ and BuOH extracts were subjected to SDS-PAGE under reducing conditions (64). Briefly, samples were dried and resuspended in 1x loading buffer (15% (v/v) glycerol, 5% (v/v) β-mercaptoethanol, 2.5% (w/v) SDS, 50 mM Tris, 1 mM EDTA, 0.0025% (w/v) bromophenol blue) and separated using 12% polyacrylamide gels, containing 0.1% (w/v) SDS, at constant 120 V. Gels were soaked in Amplify (GE Healthcare, Chicago, IL, USA), dried and exposed to films (Carestream Health Medical X-ray Blue, Rochester, NY, USA) at −70 °C. HA-tagged TbGPI2 and cMyc-tagged TbGPI1 were analyzed by immunoblotting as described before (49). EP and GPEET procyclins were analysed by immunoblotting using the LI-COR detection system (Odyssey Infrared Imager model 9120, Odyssey Application Software version 3.0.30). Primary antibodies: mouse α-GPEET 5H3 (65) at dilution 1:2500, mouse α-EP247 at dilution 1:2500. Secondary antibody: goat α-mouse IRDye 800CW at dilution 1:10000.

### Immunofluorescence Microscopy

Approximately 2.5 × 10^6^ trypanosomes were harvested by centrifugation, washed with cold PBS (137 mM NaCl, 2.7 mM KCl, 10 mM Na_2_HPO_4_, 1.76 mM KH_2_PO_4_, pH 7.4), resuspended in a small volume of PBS, spread on a microscopy slide and left to adhere for 20 min. Subsequently, parasites were fixed using 4% (w/v) paraformaldehyde in PBS for 10 min. After three washes for 5 min with cold PBS, cells were permeabilized in 0.2% (w/v) Triton X-100 for 20 min. After three additional washes and incubation in blocking solution (2% (w/v) bovine serum albumin in PBS) for 30 min, the blot was incubated with primary antibody in blocking solution for 45 min at room temperature. Antibodies used were mouse α-GPEET 5H3 (65), mouse α-EP247 (clone TRBP1/247, Cedarlane, Burlington, Canada) and mouse anti-HA (clone 16B12, Enzo Life Sciences, Farmingdale, NY, USA) at dilutions of 1:500, and rabbit anti BiP (kindly provided by J.D. Bangs, University of Buffalo, Buffalo, NY, USA) and rabbit anti-TbGRASP (kindly provided by G. Warren, Vienna Biocenter, Vienna, Austria) at concentrations of 1:2500 and 1:1500, respectively. After washing, the cells were incubated for 45 min with AlexaFluor488 or AlexaFluor594-conjugated goat α-mouse or goat α-rabbit secondary antibodies at 1:1000 dilutions in blocking buffer. After washing, the cells were mounted with Vectashield DAPI (Vector Laboratories, Burlingame, CA, USA). Immunofluorescence image stacks were captured on a Leica SP2 using a 100× oil objective. Image stacks were 3D deconvolved with the Leica LAS AF Version 2.1.0 software (Leica Microsystems CMS GmbH, Heerbrugg, Switzerland).

### Flow Cytometry

Flow cytometry was performed on live trypanosomes. Antibodies were diluted in corresponding incubation medium. Primary antibodies used were mouse α-GPEET 5H3 (65) at a dilution of 1:1000, or mouse α-EP247 at 1:500. Parasites (4 × 10^6^ cells) were harvested by centrifugation. All subsequent steps were performed at 4 °C. The cells were resuspended in 200 µl medium containing the primary antibody and incubated with rotation for 30 min. After addition of 800 µl medium, cells were pelleted and washed once with 800 µl medium. After pelleting, the cells were resuspended, incubated with Alexa Fluor 488-conjugated secondary antibodies (Invitrogen, Waltham, MA, USA, and Thermo Fisher Scientific) at dilutions of 1:1000 and washed as described for the primary antibody. After resuspension in 1.6 ml medium, fluorescence of labeled and unstained control cells was quantified with an ACEA NovoCyte benchtop flow cytometer (Agilent Technologies, Santa Clara, CA, USA). After applying a cut-off of 7.5 × 10^5^ to the forward scatter, a total of 1 × 10^4^ events were recorded and analyzed using FlowJo software (BD Life Sciences, Franklin Lakes, NY, USA) without gating.

### Social Motility

Trypanosomes were cultured in liquid medium or on semisolid agarose in a humidified incubator at 27 °C and 2.5 % CO_2_. Cells in liquid culture were maintained between 1 × 10^6^ and 1.5 × 10^7^ cells ml^−1^ by daily monitoring and diluted into the same culture flask (TPP vent screw cap, Switzerland). Semisolid culture plates containing 0.4 % (w/v) agarose/medium in a 90 mm diameter Petri dish for SoMo assays were prepared as previously described (51). Directly after air-drying for 1 h, the surface of SoMo plates was inoculated with 2 × 10^5^ cells (5 µl liquid culture concentrated to 4 × 10^7^ cells). 5 to 10 min after inoculation and without sealing the plates with Parafilm, plates were transferred to the humidified incubator. The growth pattern of trypanosome communities was documented daily with a digital camera in a dark room with LED white-light illumination from below.

### Analysis of trypanosomes cultured on SoMo plates by microscopy and propidium iodide staining

Trypanosomes were cultured on SoMo plates as described in section “Social Motility”. Cellular morphology and uptake of propidium iodide were assessed for cells grown in liquid culture or taken off SoMo plates. Samples were prepared individually, and analyzed immediately to avoid artefactual effects caused by prolonged exposure to treatments. For staining with propidium iodide, cells were taken off a SoMo plate by two washes with 1 × PBS and 300 µl of an appropriate dilution were supplemented with propidium iodide (1.0 mg/ml stock solution) to a final concentration of 5 µg/ml. Two different plates per cell line and time point were analysed. For reference, cells from liquid culture were washed twice with 1 × PBS and analysed live or after fixation (1 h in 70% EtOH on ice (66)), with and without 5 µg/ml propidium iodide. Staining with propidium iodide was quantified with a benchtop flow cytometer as described in the section “Flow cytometry”, but a cut-off of 5 × 10^5^ was applied to the forward scatter. For morphological analysis by microscopy, trypanosomes were taken off a SoMo plate 2 days post inoculation. Cells were removed from three spots by careful pipetting with 10 µl growth medium per spot. Pooled cells were paralysed by mixing 3:2 with 10% sodium azide and immediately mounted with approximately 1.5 volumes Mowiol containing 10 µg/ml Hoechst DNA stain (final concentration of approximately 1.5% sodium azide). For reference, cells from liquid culture were concentrated by centrifugation and 3 µl (∼1.5 × 10^5^ cells) were paralysed and mounted as described. Microscopy images were acquired over a period of 15 to 30 min post mounting. Differential interference contrast (DIC) and epifluorescence microscopy were performed with a Leica DM5500 instrument equipped with a DFC350 FX monochrome CCD camera using 100 × objective (HC PL APO 100X/1.40 OIL PH3 CS). Images were processed with ImageJ version 2.0.0 (Fiji).

### Cell-free labeling of GPI precursors with UDP-[^3^H]GlcNAc

Cell-free labeling was performed as previously described (30, 33). Briefly, 2.5 × 10^8^ trypanosomes were harvested by centrifugation, washed twice with PBS and hypotonically lysed on ice for 5 min in 250 µl lysis buffer (0.1 mM TLCK, 1 µg/mL leupeptin). The cell lysate was added to 250 ul HKMTLG buffer (100 mM HEPES, 50 mM KCl, 10 mM MgCl_2_, 0.1 mM TLCK, 1 µg/mL leupeptin, 20 % glycerol, pH 7.4), snap-frozen and stored at −80 °C for at least 24 h.

Before use, cell lysates were thawed, washed twice with 1 ml ice-cold HKMTL buffer (100 mM HEPES, 50 mM KCl, 10 mM MgCl_2_, 0.1 mM TLCK, 1 µg/mL leupeptin, pH 7.4), resuspended in 150 µl 2x HKMTL buffer containing 10 mM MnCl_2_ and 0.2 µg/ml tunicamycin and pre-warmed to 27 °C. For each sample, 1 µl (equal to 1 µCi) UDP-[^3^H]GlcNAc was added to 30 µl pre-warmed DA buffer (2 mM DTT, 2 mM ATP). 30 µl of cell lysate was added and the mixture was incubated at 27 °C for the indicated time. The assay was terminated by addition of 400 µl chloroform/methanol 1:1 (v/v) for a final chloroform/methanol/water ratio of 10:10:3 (v/v/v). GPI precursors were solubilized by water bath sonication and incubation on ice. Insoluble compounds were removed by centrifugation. The pellet was re-extracted with 250 µl chloroform/methanol/water 10:10:3 (v/v/v). The extracts were pooled, dried under N_2_ and purified by partitioning between n-butanol and water. The upper (organic) phase was collected while the lower phase was re-extracted with water-saturated butanol, both organic phases were pooled and remaining impurities were removed by back-extraction with butanol-saturated water. The purified organic phases were subjected to β-counting and TLC analysis as described above.

### Mass spectrometry analysis of procyclin-derived GPI anchors

Extraction and purification of procyclins. Procyclins (both GPEET and EP forms) were purified from 10^10^ cells by organic solvent extraction and octyl-Sepharose chromatography as previously described (27, 28) but with a slight modification. Briefly, the cells were extracted three times with chloroform/methanol/water (10:10:3, v/v). After the delipidation process, the pellet was dried under N_2_ and subsequently extracted twice with 9% butan-1-ol in water. The supernatant of 9% butan-1-ol extracts were pooled and dried under N_2_. To further purify the extracted procyclins, the dried samples were redissolved in 1 ml of buffer A (5% propan-1-ol in 0.1 M ammonium acetate) and applied to 0.5 ml of octyl-Sepharose 4B packed in a disposable column and pre-equilibrated with buffer A. The column was washed with 3 ml of buffer A followed by 3 ml of buffer B (5% propan-1-ol). The procyclins were eluted in 2.5 ml of buffer C (50% propan-1-ol) and concentrated and dried by rotary evaporation.

Permethylation and ES-MS of GPI glycans. The dried procyclin samples were treated with 100 µl of ice-cold 50% aqueous hydrogen fluoride (aq. HF) for 24 h at 0 °C to cleave the GPI anchor ethanolamine-phosphate-mannose and inositol-phosphate-acylglycerol phosphodiester bonds. The samples were freeze dried to evaporate the remaining aq. HF and redissolved in 100 µl water and centrifuged at 16000 × g for 10 min. The supernatant containing the GPI glycans was taken for permethylation by the the sodium hydroxide method, as described (7, 8). The permethylated GPI glycans, bearing a fixed positive charge in the form of an *N*-trimethyl-glucosamine quaternary ammonium ion, were dissolved in 100 µl of 80% acetonitrile, and 10% aliquots were dried and recovered in 10 µl of 80% acetonitrile, 0.5 mM sodium acetate. The samples were infused into the Orbitrap Fusion tribrid mass spectrometer (Thermo Scientific) using static infusion nanoflow probe tips (M956232AD1-S, Waters). Data were collected in positive ion mode for ES-MS, ES-MS^2^ and ES-MS^3^. Positive ion spray voltage was 0.7 kV and the ion transfer tube temperature was 275 °C. Collision induced dissociation (CID) was used for MS^2^ and MS^3^ fragmentation, using 25-35% collision energy.

### Protein analysis by native PAGE

The GPI GlcNAc transferase complex was immunoprecipitated from parasites expressing myc-tagged TbGPI1 as previously described (39). Briefly, 2 × 10^8^ trypanosomes were harvested by centrifugation, washed twice with PBS and lysed for 30 min on ice in 500 µl lysis buffer (50 mM Tris-HCl, 150 mM NaCl, 1% digitonin, pH 7.4). Insoluble components were removed by centrifugation at 16000 g for 20 min at 4 °C. The supernatant was incubated with α-cMyc agarose beads (Takara Bio, Kyoto, Japan) on a rotary wheel for 16 h at 4 °C. Bound complexes were eluted three times with 10 μL lysis buffer containing 0.5 mg/mL cMyc peptide. Eluated complexes were subjected to native PAGE as previously described (67). Briefly the protein complexes were supplemented with 10× BN loading buffer (5% (w/v) Coomassie Brilliant Blue G-250, 500 mM 6-aminocaproic acid, 100 mM Bis-Tris·HCl, pH 7.0) and separated on a 4-15% polyacrylamide gel (Bio-Rad Laboratories, Hercules, CA, USA). After separation, proteins were transferred on a polyvinylidene fluoride membrane and detected by immunoblotting against the cMyc epitope.

## Notes

* The work was supported by Swiss National Science Foundation Sinergia grant CRSII5_170923 to RH, AKM and PB, by a Wellcome Tust Investigator Award (101842/Z13/Z) to MAJF and Swiss National Science Foundation grant 310030_184669 to IR. The authors declare that they have no conflicts of interest with the contents of this article.

